# Self-Other Neural Representational Distinction is Associated with Distorted Social Predictions in Social Anxiety

**DOI:** 10.64898/2026.10.29.762906

**Authors:** Yanyan Qi, Yun Chen, Yixiao Hu, Qianqian Fan, Lingxiao Wang, Guochun Yang, Grit Hein, Haiyan Wu

## Abstract

**Background:** Social anxiety (SA) is characterized by distorted negative expectations about the future. Because optimism bias is inherently comparative, the extent to which these predictions reflect a fundamental disruption in self- versus other-referential processing remains poorly understood.

**Methods:** Across two experiments, we investigated the mechanisms underlying social predictions in SA. In Experiment 1, individuals with high (*n* = 52, LSAS > 65) and low SA (*n* = 51, LSAS < 35) rated the future likelihood of positive and negative social outcomes for themselves or for a protagonist. In Experiment 2, 35 adults spanning a continuous spectrum of SA completed the task during fMRI. Behavior was analyzed using linear mixed-effects models. Neurally, multivariate pattern analysis (MVPA) quantified self-other representational distinctions, while a parametric general linear model (GLM) tracked neural sensitivity to likelihood predictions.

**Results:** Behaviorally, higher SA was characterized by an attenuated optimism bias: individuals predicted a lower likelihood of positive outcomes, but a higher likelihood of negative outcomes, for themselves compared to others. Neurally, higher trait SA correlated with greater self-other representational distinction within the left pregenual anterior cingulate cortex (pgACC). Concurrently, the precuneus exhibited stronger parametric tracking of self-negative predictions versus others. Crucially, pgACC representational distinction during positive outcome processing significantly mediated the association between SA severity and heightened prediction-tracking in the precuneus.

**Conclusions:** These findings offer a novel mechanistic framework, suggesting that an exaggerated neural segregation of the self from others drives hyper-vigilant, risk-sensitive social predictions in SA. This candidate circuit-level dynamic warrants further replication in clinical cohorts.

## Introduction

Prediction is fundamental to adaptive human existence. Individuals typically exhibit a robust optimism bias—more precisely, comparative (unrealistic) optimism—characterized by the tendency to overestimate the likelihood of positive events occurring to oneself relative to others, while simultaneously underestimating the risk of negative outcomes (1–3). This bias serves as a vital psychological shield, maintaining self-esteem and mitigating anticipatory anxiety within an uncertain social landscape. However, this protective mechanism erodes significantly along the dimensions of anxiety and depression (4,5). In particular, socially anxious individuals may exhibit a diminished optimism bias in social contexts, since they often hold negative expectations during social interactions (6–8). Yet, the neural mechanisms that determine how this protective shield is compromised in relation to social anxiety (SA) remain poorly understood.

Existing studies on optimism bias have primarily focused on the cognitive mechanism of self, treating individuals as the central unit of analysis in self-referential imagery (9–13) or reactions to personal prediction errors (2,14–16). While these studies incorporate social information, they rarely examine the neural distinction between self and other, even though optimism bias is fundamentally a comparative judgement between these two entities (3). Therefore, understanding why this bias fails in SA requires a shift in perspective toward how the brain differentially processes self- and other-referential information.

Decades of neuroimaging research have established that the processing of self and others engages a consistent “social brain” network, primarily involving the medial prefrontal cortex (mPFC), precuneus, and key nodes of the theory-of-mind circuit, such as the temporoparietal junction (TPJ) and superior temporal sulcus (STS) (17–21). While early studies primarily compared the average activation levels evoked by self-versus other-referential processing, recent advances using multivariate pattern analysis (MVPA) have shifted the focus toward the fine-grained spatial patterns of these responses. These patterns reveal a representational structure where social closeness is encoded as neural similarity: the brain represents those we feel close to in a manner strikingly similar to how it represents the self (22–24). This suggests that the brain maps the social world onto a continuum of self-similarity. However, it remains unknown whether this neural similarity between self and other provides the underlying neural foundation for the predictions we make for ourselves versus others.

The neural similarity is thought to facilitate self-projection, a process where the brain utilizes self-referential templates to simulate and understand the experiences of others (22,24–27). However, this adaptive mapping may be fundamentally compromised in SA. Characterized by an intense fear of negative evaluation and a profound sense of social aloneness, SA is often accompanied by hyper-heightened self-focused attention, where individuals become excessively preoccupied with their internal states and perceived social flaws (8,28–32), which may be in turn reflected by an exaggerated neural segregation between the self and others. Instead of the adaptive neural overlap seen in healthy individuals (22,33–35), the socially anxious brain may partition the self as an isolated target. This increased representational distinction (i.e., lower neural similarity) may act as a computational constraint, biasing the neural tracking of social risk during the predictive process regarding oneself versus others.

To test this mechanism, we conducted two complementary experiments using a validated social prediction paradigm (Figure 1). Building on evidence that socially anxious individuals manifest a coexistence of fear concerning both negative and positive evaluations (36,37), the present study required participants to imagine both positive and negative social events (e.g., receiving applause or being denied help; Table S1) occurring to either themselves or a gender-neutral other (“Liu Yang”). Experiment 1 utilized a behavioral cohort (high versus low SA) to establish the attenuated optimism bias. Experiment 2 employed functional magnetic resonance imaging (fMRI) in a continuous SA cohort to uncover the underlying neural mechanisms.

**Figure 1.**
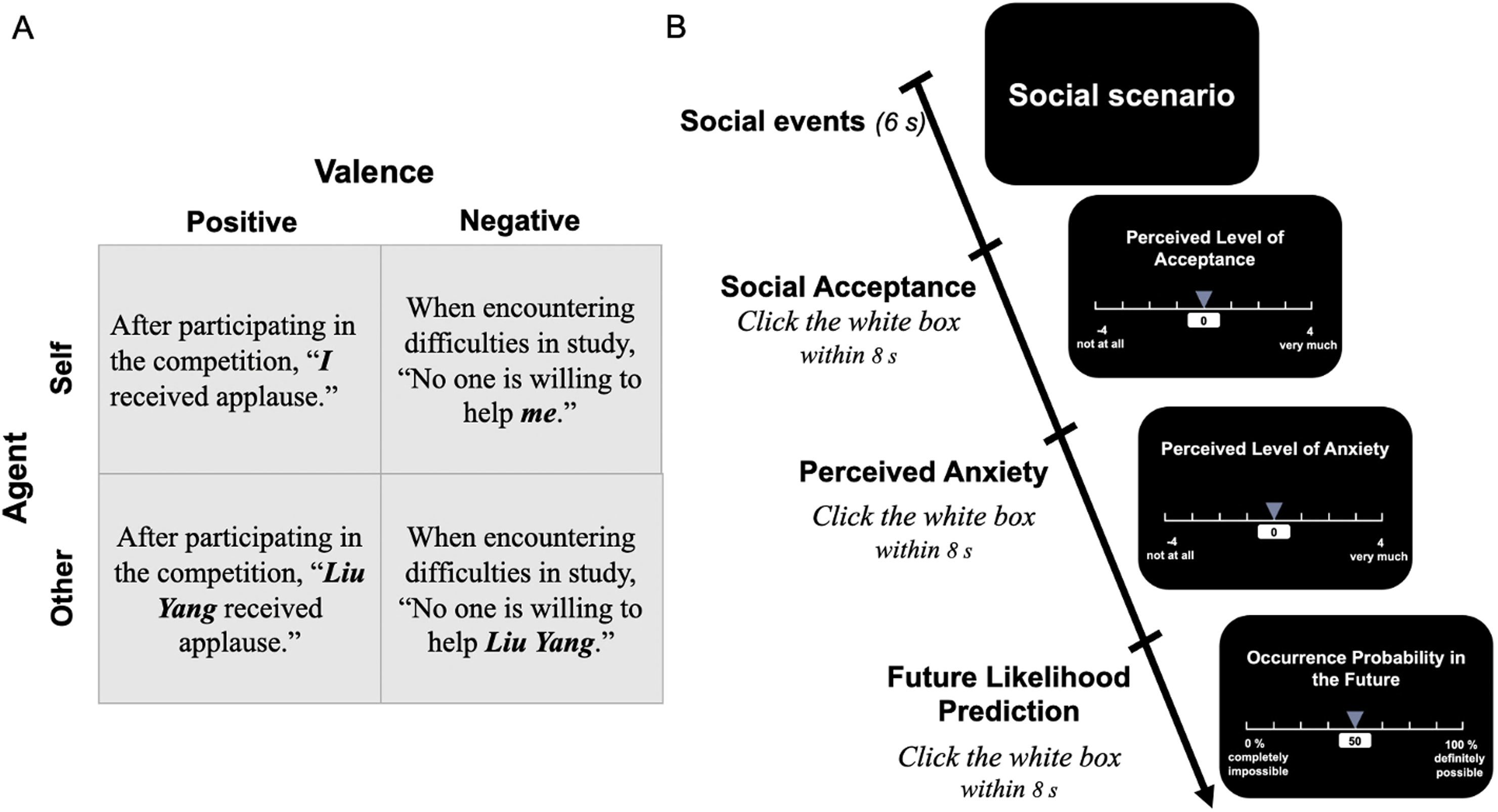
Experimental Design and Trial Structure. (A) Experimental Factors. The study employed a 2×2 within-subject design with Agent (Self/ Other) and Valence (Positive/ Negative) as factors. Example social scenarios are shown for each of the four experimental conditions. **(B) Timeline of a Single Trial**. A standard trial consisted of a social event presentation (6 s), followed by three distinct rating phases, each with an 8 s time limit: Perceived Social Acceptance, Perceived Anxiety, and Likelihood Prediction for future occurrences.

By integrating MVPA decoding to quantify self-other neural representational distinction with parametric modulation to identify the neural correlates of trial-by-trial social prediction, we focused on key Regions of Interest (ROIs, Figure S1) within the broader social brain network. These included core mentalizing and self-referential hubs, such as the mPFC, precuneus, left and right TPJ, and left and right STS (17–21,38), as well as regions critical for monitoring personal relevance and emotional salience (e.g., social acceptance and rejection), specifically the left and right pregenual anterior cingulate cortex (pgACC) and left and right anterior insula (AI) (14,39–42). Our focus on these specific nodes is directly supported by recent evidence: a systematic review reports heightened medial prefrontal-anterior cingulate-insula engagement during self-referential processing in SA (43), and resting-state work shows that precuneus-perigenual ACC connectivity scales inversely with SA severity along a dimensional continuum (44). This integrated approach allowed us to examine whether SA biases social prediction through a fundamental segregation of the self from others across both cognitive and affective neural manifolds.

We tested three interconnected hypotheses. First, at the behavioral level (Experiments 1 and 2), we hypothesized that higher SA would manifest as an attenuated self-referential optimism bias. Second, neurally, higher SA would be associated with an increased neural segregation (i.e., higher decoding accuracy) between self and other. Finally, this representational segregation may serve as an underlying mechanism that drives the divergent computational tracking of social predictions for oneself versus others. Specifically, we tested an exploratory mediation model investigating that self-other neural segregation accounts for the association between between SA and altered prediction-tracking sensitivity. Taken together, our three hypotheses outline a directional cascade in which SA first amplifies the neural segregation of the self from others, and this representational isolation, in turn, biases the neural tracking of self-relevant social threat more vigilantly than other-relevant prediction.

## Results

### Behavioral Evidence of Reduced Optimism Bias in SA

#### Experiment 1: Categorical Comparison of High and Low SA

In Experiment 1, the linear mixed model (LMM) with SA (high versus low), Valence (positive versus negative), Agent (self versus other), and their interaction as independent variables revealed significant main effects of Valence (χ*²*(1) = 4053.091, *p* < 0.001) and Agent (χ*²*(1) = 10.295, *p* = 0.001); alongside a three-way interaction of SA × Valence × Agent (χ*²*(1) = 102.724, *p* < 0.001; Figure 2A; Table S2). Further analysis showed that for positive events, the interaction of SA × Agent was significant (χ*²*(1) = 67.456, *p* < 0.001; Figure 2A; Table S2); the low SA group estimated comparable possibility of positive events occurrence for oneself (*M* = 67.717, *SE* = 1.740) and for others (*M* = 69.936, *SE* = 1.740; *p* = 0.748). In contrast, the high SA group estimated a significantly lower possibility for oneself (*M* = 59.665, *SE* = 1.719) compared to others (*M* = 67.773, *SE* = 1.719; *p* < 0.001). For negative events, the SA × Agent was significant (χ*²*(1) = 52.183, *p* < 0.001; Figure 2A; Table S2); low SA group estimated a significant lower possibility for oneself (*M* = 40.023, *SE* = 2.033) than for other (*M* = 42.001, *SE* = 2.033; *p* = 0.0068), while the high SA group estimated a significantly higher possibility for oneself (*M* = 45.207, *SE* = 2.033) than for others (*M* = 39.753, *SE* = 2.009; *p* < 0.001). These findings showed an attenuated optimism bias in individuals with high SA.

**Figure 2.**
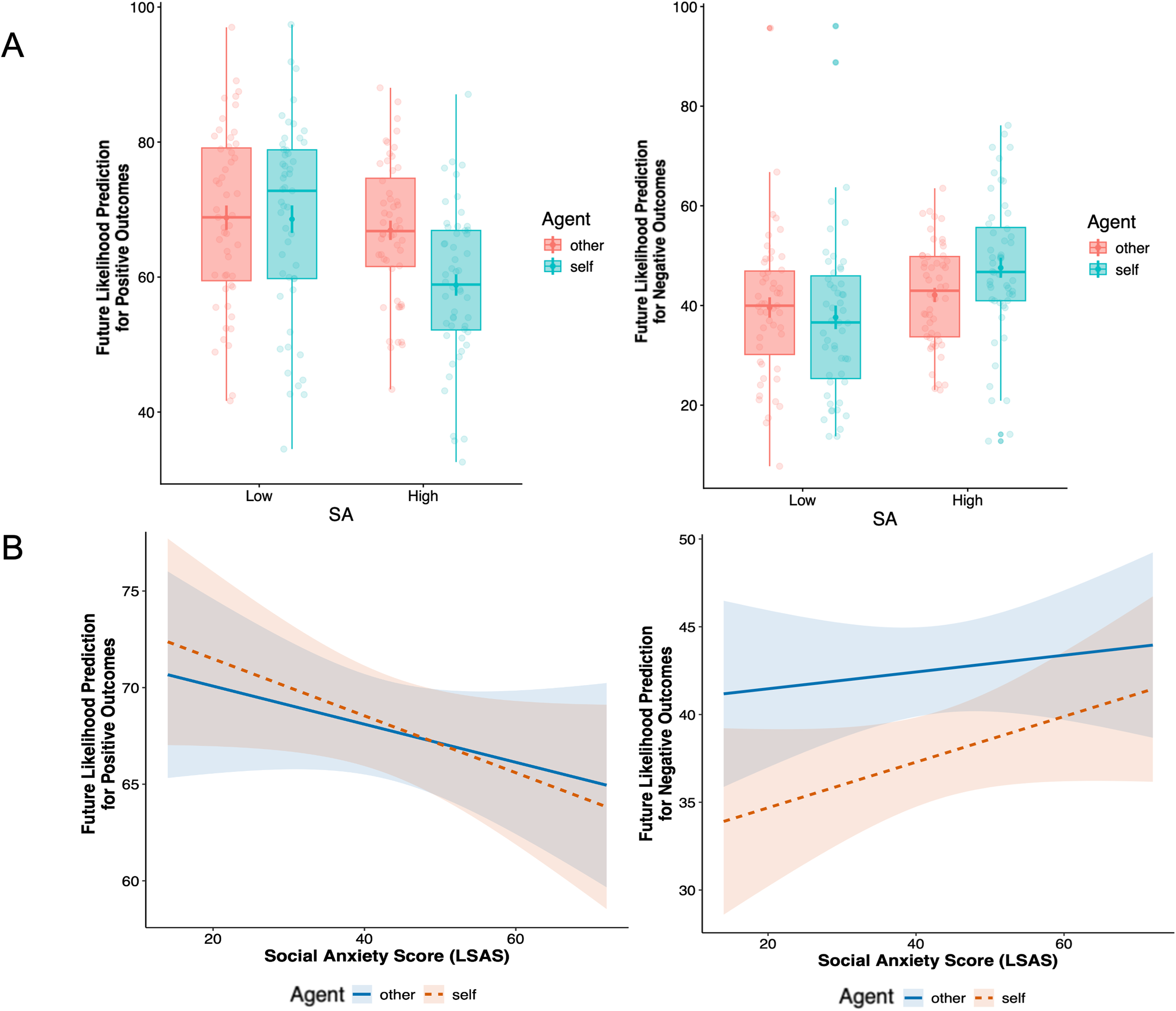
Behavioral Results. (A) Experiment 1. Predicted likelihood of future positive (left) and negative (right) outcomes for individuals with low and high social anxiety (SA), separately for self- and other-referential judgments.**(B) Experiment 2**. Predicted likelihood of future positive (left) and negative (right) outcomes as a function of continuous social anxiety symptoms, indexed by the Liebowitz Social Anxiety Scale (LSAS), separately for self- and other-referential judgments. The shaded area indicates the 95% confidence interval.

#### Experiment 2: Replication with Continuous SA

In Experiment 2 (continuous SA), results replicated these trends. The LMM revealed a significant three-way interaction among Agent, Valence, and SA (χ*²*(1) = 4.198, *p* = 0.040; Figure 2B; Table S3). To decompose three-way interaction effect, we conducted follow-up LMMs separately for the positive and negative conditions. In the positive condition, no significant interaction between Agent and SA was observed (χ*²*(1) = 1.645, *p* = 0.200; Figure 2B; Table S3). However, in the negative condition, a trend-level interaction between Agent and SA was observed (χ*²*(1) = 3.417, *p* = 0.065; Figure 2B; Table S3). Simple slope analysis revealed that while the predicted occurrence likelihood of negative outcomes for others remained relatively stable across the SA level (*b* = 0.051, *p* = 0.6128), the association between SA and the predicted occurrence likelihood for the self showed a more positive trend (*b* = 0.135, *p* = 0.192; Figure 2B). Although neither simple slope reached statistical significance; the divergence between the two slopes is directionally consistent with the three-way interaction of Experiment 1. **Brain Findings**

#### Higher SA is Associated with Stronger Neural Differentiation Between Self and Other

ROI-based decoding analyses showed that in the positive valence condition, neural patterns in right TPJ (*M* = 0.553, 95% CI lower bound = 0.522, *p*_FDR_ = 0.034) significantly discriminated between self and other (Figure 3A; Table S4). In the negative valence condition, successful decoding of self versus other was observed in the left STS (*M* = 0.565, 95% CI lower bound = 0.532, *p*_FDR_ = 0.025; Figure 3B; Table S4).

**Figure 3.**
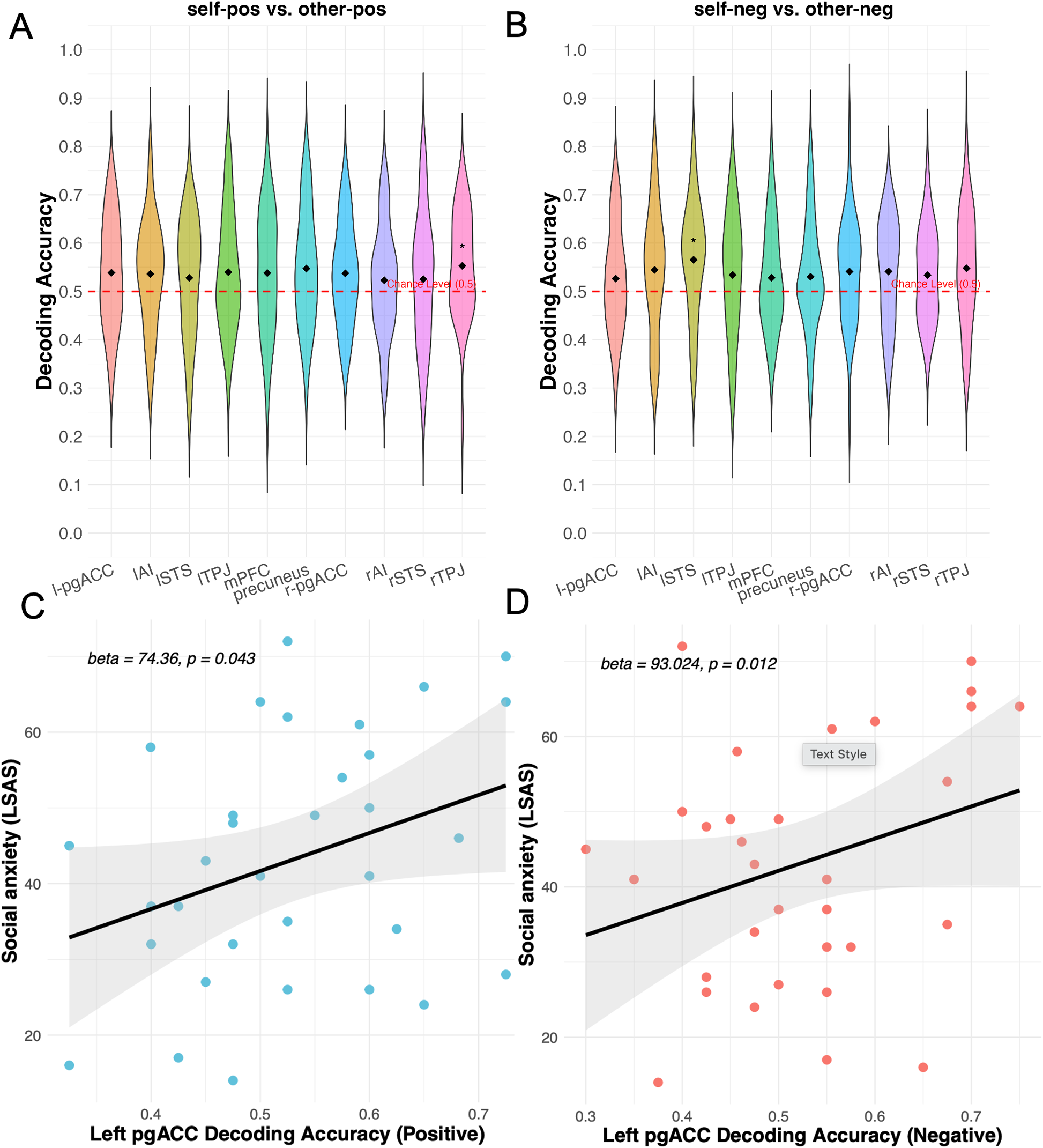
Multivariate Decoding of Self versus Other and Its Correlation with Social Anxiety (SA). (A–B) Regional Decoding Accuracies for Self versus Other. Violin plots illustrate the decoding performance above chance level (0.5, red dash line) across various regions of interest (ROIs) in (A) positive and (B) negative social contexts. * *p_FDR_* < .05. **(C–D) The Positive Association between Decoding Accuracy of pgACC and SA (LSAS total score) in (C) Positive and (D) Negative Conditions.** The displayed regression lines are bivariate fits for visualization purposes; the reported coefficients derive from multiple regression models including all 10 ROIs as predictors (Tables S5 and S6). Solid lines represent linear fits with 95% confidence intervals (shaded areas). l-/ r-pgACC: left/ right pregenual anterior cingulate cortex; lSTS/ rSTS: left/ right superior temporal sulcus; lTPJ/ rTPJ: left/ right temporoparietal junction; mPFC: medial prefrontal cortex; lAI/ rAI: left/ right anterior insula.

Next, a multiple linear regression model in the positive condition revealed that decoding accuracy in the left pgACC emerged as the strongest predictor among all the ROIs, showing a significant positive association with SA levels (*b* = 74.364, *p* = 0.043*; R*² = 0.370; Figure 3C; Table S5). A similar pattern was observed in the negative condition, where the left pgACC again demonstrated the strongest predictive effect (*b*= 93.024, *p* = 0.012; *R*^2^ = 0.473; Figure 3D; Table S6). These results highlight a pivotal role for the left pgACC. Of note, left pgACC decoding accuracy did not significantly exceed chance at the group level after FDR correction in either condition (positive: *p*_FDR_ = 0.051; negative: *p*_FDR_ = 0.098; Tables S5 and S6); the results reported here therefore reflect individual-difference associations with decoding scores rather than group-level significant decoding.

To complement our multivariate approach, exploratory whole-brain univariate analyses were conducted to examine both general self-versus other-referential differences and their modulation by SA severity. These analyses revealed a normative self-positivity bias (11,45) and yielded individual-difference patterns consistent with our MVPA results (e.g., SA positively correlating with mPFC/pgACC hyperactivation). Full details are provided in the Supplementary Results and Tables S7–S9.

#### Selective Neural Tracking of Negative rather than Positive Social Outcome Prediction

The parametric GLM analysis revealed a distributed neural network significantly tracking trial-by-trial subjective likelihood ratings for self-negative social outcomes. Specifically, brain regions (Table 1) including bilateral STS, precuneus, and right pgACC, as well as the bilateral inferior frontal gyrus (IFG), showed a linear increase in response as the perceived probability of negative social outcomes for the self rose. In contrast, no significant clusters were identified for self-positive, other-positive or other-negative predictions, nor did any regions significantly track perceived anxiety or acceptance ratings after FWE correction. d *t*-tests directly comparing self-versus other-negative tracking isolated a significant effect exclusively in the right precuneus (*p*_FWE_ = 0.005; Figure 4A), with no significant activations observed in other conditions or dimensions. Given the precuneus’s established role in self-referential mental imagery and first-person perspective-taking (17,46,47), this heightened activation suggests a more intensive neural engagement when simulating negative social futures for the self compared to others.

**Figure 4.**
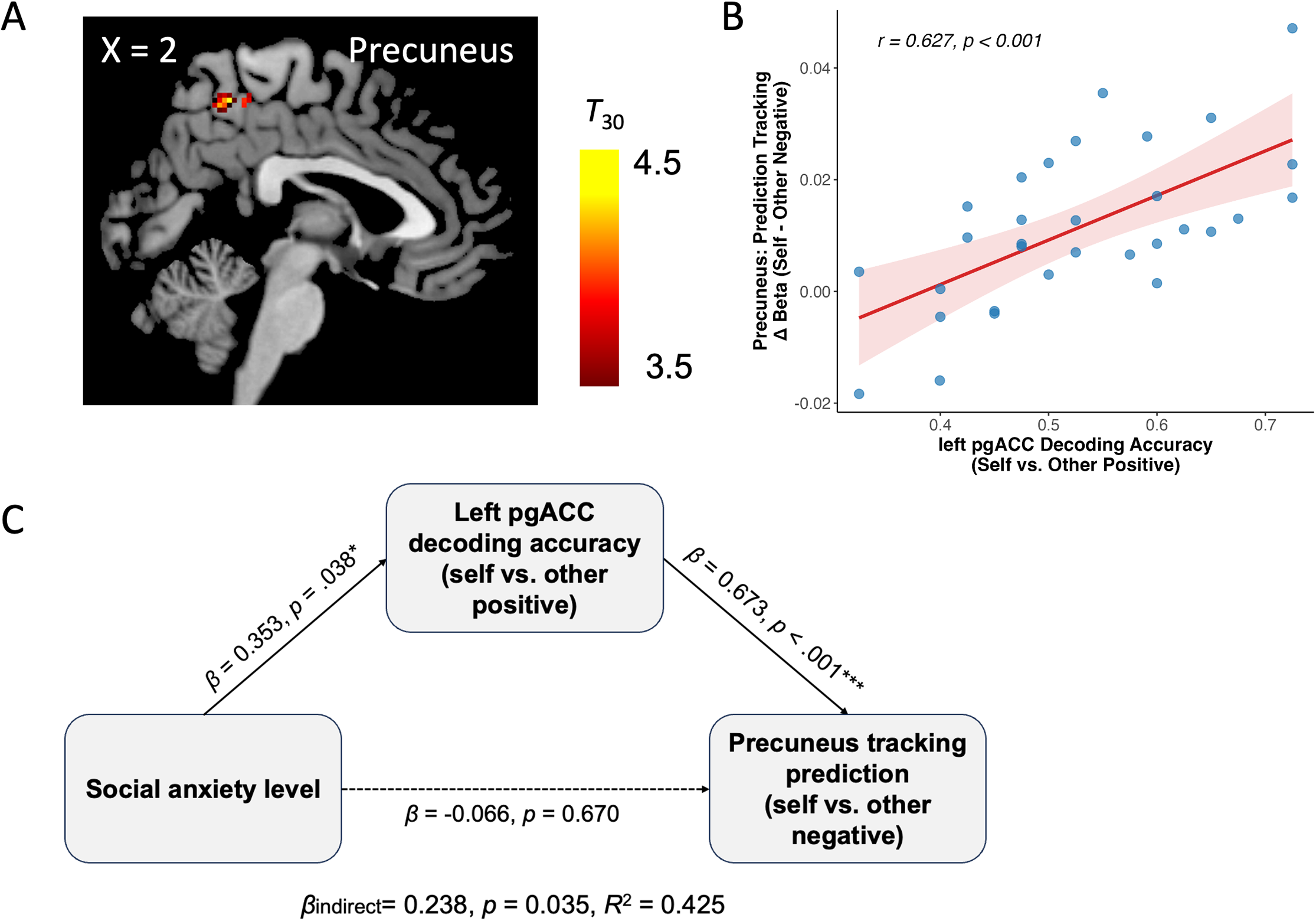
Precuneus Neural Tracking and Its Mediation by Pregenual Anterior Cingulate Cortex (pgACC) Decoding. (A) Neural Tracking of Social Prediction for Self versus Other in Precuneus. The sagittal slice illustrates the significant cluster in the right precuneus identified during the neural tracking of negative social predictions (self versus other). (B) **Spearman Correlation between Decoding Accuracy and Prediction Tracking.** A significant positive correlation was found between the left pgACC decoding accuracy (self versus other positive) and the precuneus tracking sensitivity (Δβ: self-negative versus other-negative prediction; *r* = 0.627, *p*_uncorrected_ < .001, *p*_FDR_ = .003). **(C) Exploratory Mediation Model**: the left pgACC decoding accuracy (self versus other positive) showed a significant indirect association linking SA levels and precuneus tracking of social predictions (self versus other negative). * *p* < .05, *** *p* < .001.

**Table 1.**
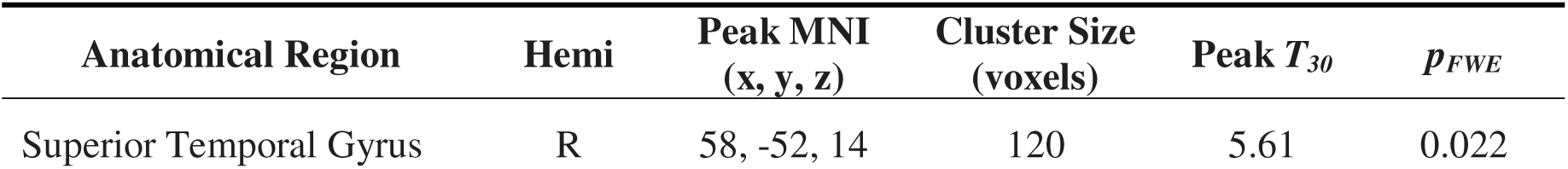
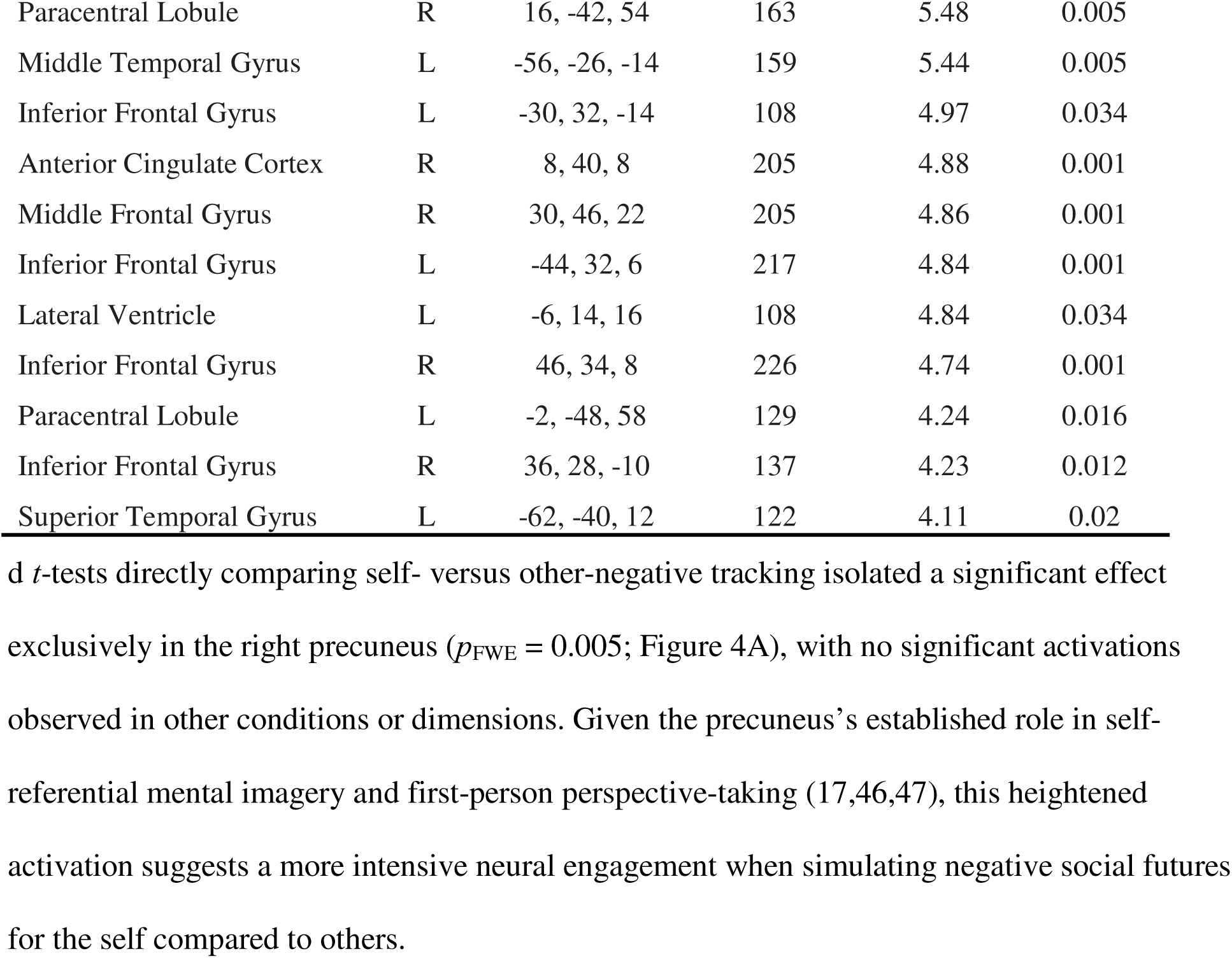
Brain regions showing significantly trial-by-trial tracking of future likelihood for negative social outcomes.

#### Left pgACC Representational Distinction Correlates with Precuneus Predictive Tracking

Spearman correlations revealed a robust positive correlation between the decoding accuracy of the left pgACC (self-positive versus other-positive) and the neural sensitivity of the precuneus during self-negative versus other-negative prediction (*r* = 0.627, *p*_uncorrected_ < 0.001, *p*_FDR_ = 0.003; Figure 4B; Table S10). This association suggests that the self-versus-other neural representational distinction when processing positive social outcomes in the left pgACC is intrinsically linked to how the precuneus computationally differentiates between self and other when forecasting negative social outcomes. Two additional pairs involving mPFC and right TPJ also showed marginal associations during the positive condition (both *p*_FDR_ = 0.079; Table S10), suggesting that this pattern may not be entirely exclusive to the pgACC. Notably, however, the pgACC–precuneus pair was the only one among the 20 tested ROI–valence pairs to survive FDR correction and was thus selected for the following mediation analysis.

#### The Mediation Model

To synthesize our findings, we constructed a mediation model using a path-analytic framework. The mediation analysis (Figure 4C) revealed a significant indirect effect of SA on precuneus prediction sensitivity via left pgACC representational distinction (indirect effect β = 0.238, bias-corrected bootstrap 95% CI = [0.016, 0.459], 5,000 resamples). Higher SA was significantly associated with enhanced neural differentiation between self and other within the left pgACC (path *a*: β = 0.353, 95% CI = [0.020, 0.687], *p* = 0.038). This intensified representational distinction, in turn, strongly predicted a higher neural sensitivity to social prediction signals in the precuneus (path *b*: β = 0.673, 95% CI = [0.439, 0.907], *p* < 0.001). Crucially, the direct effect of SA on precuneus sensitivity became non-significant after accounting for the mediator (path *c*′: β = −0.066, 95% CI = [−0.371, 0.239], *p* = 0.670). This pattern indicates a significant indirect effect alongside a non-significant direct path.

## Discussion

In the present study, we investigated the behavioral and neural mechanisms underlying distorted social predictions in SA, specifically examining the role of self-versus other-referential processing. Behaviorally, our two complementary experiments revealed that higher SA is characterized by a significantly attenuated optimism bias, where individuals anticipate fewer positive and more negative social outcomes for themselves relative to others. Neurally, we identified that SA is associated with an exaggerated representational segregation between the self and others within the pgACC. Furthermore, this pgACC representational distinction mediated the relationship between SA severity and hyper-vigilant tracking of negative social predictions in the precuneus. Together, these findings suggested that disrupted neural integration of self-other representations drives the negative social expectations observed in SA.

### Reconciling attenuated optimism bias with the anxiety literature

Both experiments demonstrated that SA is characterized by an eroded optimism bias. This pattern should be considered alongside the findings of (4), who reported that reduced optimism was specific to generalized anxiety disorder and was not observed in SA disorder. The two studies differ in several respects—dimensionally assessed trait SA in a non-clinical sample versus diagnostic patient groups, socially evaluative vignettes versus everyday events—so their results are not directly comparable. One reconciling possibility is that attenuated optimism in SA is specific to socially evaluative contexts, whereas the optimism reduction in generalized anxiety generalizes across contexts; testing this hypothesis will require clinical samples assessed across both content domains.

### Neural segregation of self and other in the pgACC

Neurally, the MVPA focused on key nodes within the ToM and salience networks. While several regions exhibited overlapping patterns for self and other, specific areas including rTPJ and lSTS showed decoding accuracies significantly above chance (0.5) (see Figure 3). These results suggest a nuanced neural architecture that supports both shared and distinct representations of social agents, consistent with previous models of self-other integration (33,35).

Crucially, regression analyses revealed that heightened SA was strongly associated with increased left pgACC decoding accuracy across both positive and negative conditions. The pgACC is a critical hub for integrating affect and self-referential information, particularly in monitoring social harmony and positive self-regard, such as social acceptance (39,42,45). The intensified differentiation observed in the pgACC during positive outcomes may reflect a vigilant separation of the self from others when facing positive social feedback. Conversely, the heightened distinctness in negative condition suggests that SA may be accompanied by a more rigid and intense monitoring of the self when facing social threats. This interpretation is firmly supported by recent systematic evidence highlighting hyper-engagement of the anterior cingulate cortex and mPFC during self-referential processing as a core neural signature of social anxiety (43). In socially anxious individuals, this representational sharpening may be a neural manifestation of the fear of both positive and negative evaluations (36,37). These findings align well with clinical accounts of self-focused attention in SA (29–32), where the neural representation of the self becomes selectively sharpened against the other under perceived social risk.

### Self-other neural similarity and the “lonely mind” of SA

Furthermore, in light of recent evidence that interpersonal closeness is typically tied to increased self-other neural similarity (22–24), our finding SA severity associates positively with neural dissimilarity (i.e., higher decoding accuracy) indicates that the “lonely mind” of the socially anxious individual is characterized by a pronounced neural dissimilarity across both positive and negative social evaluations.

### Self-other predictive tracking in the precuneus

Regarding predictive tracking, a network comprising the mPFC, STS, IFG, and ACC (see Table 1), regions associated with self-referential processing, cognitive control, and salience monitoring (Apps et al., 2016; Li et al., 2017; Menon, 2023; Schurz et al., 2014; van Veluw & Chance, 2014), dynamically tracks one’s estimated likelihood of negative social events. Crucially, when comparing self-versus other-related predictions, the precuneus exhibited significantly stronger tracking for self-negative likelihoods. The precuneus is not only a core hub of DMN which is robustly associated with resting-state self-reflection, but is also actively engaged in cognitive tasks related to reward/ risk estimation (Dadario & Sughrue, 2023; Utevsky et al., 2014). The prioritized tracking observed here suggests that the brain generally allocates more computational resources to self-relevant negative outcomes, reflecting a fundamental self-centered bias in social anticipation (Beer, 2014; Hughes & Beer, 2013).

Synthesizing these mechanisms, our mediation model demonstrated that increased pgACC representational distinction during positive outcomes in higher SA exacerbates the biased precuneus tracking of self-negative predictions. The functional interplay between these two specific nodes is corroborated by recent resting-state evidence demonstrating that intrinsic precuneus-pgACC connectivity scales inversely with social anxiety severity along a dimensional continuum (44), providing an independent network-level basis for our task-based mediation findings. Although the cross-valence nature of this finding might initially appear counterintuitive, it is highly consistent with previous behavioral evidence by Alden et al. (2008). They demonstrated that socially anxious individuals who interpret positive evaluation through a negative lens tend to hold significantly more pessimistic social predictions. Our findings extend this observation to the neural level: representational isolation in the pgACC reflects a failure to process positive social value similarly with others, which may potentially leave the individual more vulnerable to the hyper-vigilant self versus other neural tracking of social threat prediction in the precuneus. This intersects with recent work isolating the neurocognitive signatures of fear of positive evaluation (36), which further motivates treating positive and negative evaluation as co-active threats in SA. Taken together, this cross-valence cascade provides a novel mechanistic account of how the fear of positive evaluation and fear of negative evaluation are neurally intertwined in SA.

### Limitations and Translational Implications

Despite these novel insights, several limitations of the present study should be acknowledged. First, although Experiment 1 utilized a well-powered behavioral cohort, the underlying neural mechanisms and the subsequent mediation model were derived from the more modest fMRI sample in Experiment 2 (*n* = 31–35). Consequently, the statistical power for our individual-difference neuroimaging analyses is constrained, rendering the mediation results preliminary and hypothesis-generating. Furthermore, because both experiments relied on non-clinical samples to assess dimensional trait SA, it remains to be determined whether these patterns fully extend to diagnosed social anxiety disorder (SAD). Second, the cross-sectional nature of our design precludes definitive causal inferences. While the pgACC–precuneus mediation model is well grounded, alternative causal structures (e.g., reversed or common-cause models) cannot be excluded. Future longitudinal studies in clinical cohorts are necessary to establish the causal dynamics of this neural pathway and verify its relevance to diagnosed SAD.

If the pgACC-mediated self-other segregation that drives hyper-vigilant precuneus tracking generalizes to clinically diagnosed SAD, it nominates a candidate circuit-level target for mechanism-focused interventions. Recent computational-psychiatry accounts of SAD (48) emphasize that biased social learning and self-referential processing are tractable intervention targets; accordingly, interventions that bridge the neural self-other gap-for example, oxytocin administration (26), social-cognition training, or neurofeedback aimed at increasing pgACC self-other overlap-may help restore the representational integration required for healthy social functioning. Such predictions should be tested in pre-registered clinical trials with adequate statistical power in future studies.

In conclusion, this study demonstrated that SA is associated with distorted social predictions. Neurally, higher trait SA is associated with a pronounced neural representational sharpening within the pgACC, reflecting a reduced tendency to process one’s positive social feedback similarly to others. This segregation is further coupled with hyper-vigilant tracking of self-versus other-referential social threat predictions in the precuneus. If replicated longitudinally and in clinical samples, this cross-valence neural pattern could inform mechanism-focused interventions for SA, for example, by testing whether interventions that bridge the neural self-other gap help restore the representational integration necessary for healthy social functioning.

## Materials and Methods

### Participants

Across both experiments, all participants were right-handed, had normal or corrected-to-normal vision, reported no history of psychiatric or neurological disorders, and provided written informed consent. The protocols complied with the Declaration of Helsinki and were approved by the Institutional Review Board at the University of Macau (BSERE21-APP005-ICI).

For Experiment 1, based on a priori power analyses (see Supplementary Methods), we recruited 52 participants for the high SA group (Liebowitz Social Anxiety Scale (LSAS; 49) > 65; *M* = 21.50 years, *SD* = 2.06; 29 females) and 51 participants for the low SA group (LSAS < 35; *M* = 21.20, *SD* = 2.15; 30 females). For Experiment 2, 36 adults spanning a continuous spectrum of trait SA were initially recruited (*M* = 22.90, *SD* = 3.50; 25 females). Following data collection, one individual was excluded due to excessive head motion, leaving 35 participants for the MVPA. Four additional participants were excluded from the parametric GLM and mediation analyses due to zero variance in behavioral ratings, resulting in a final sample of 31 participants.

### Stimuli and Experimental Design

Both experiments utilized a social prediction paradigm manipulating Agent (self versus other) and Valence (positive versus negative) (Figure 1). In Experiment 1, this was structured as a 2 × 2 × 2 mixed-model design, incorporating SA (high versus low) as a between-subjects factor. In Experiment 2, the task was adapted into a 2 × 2 within-subjects factorial design to assess continuous SA individual differences. During task, participants mentally simulated social scenarios occurring either to themselves or a protagonist, and subsequently rated perceived social acceptance, perceived anxiety, and future likelihood prediction—dimensions explicitly chosen because perceived unacceptance and automatic anxiety are core features of social anxiety disorder (American Psychiatric Association, 2013).

In Experiment 1, participants completed the full set of 54 social scenarios: 27 positive (e.g., “Receiving a warm ovation after a public speech”) and 27 negative (e.g., “Facing a lack of support during academic challenges”), yielding 108 trials. In Experiment 2, to minimize participant fatigue during fMRI scanning, the task was optimized after the first four participants (who completed the original 108-trial version). The remaining 32 participants completed an 80-trial version using a representative subset of 40 scenarios. Both versions were rigorously balanced for scenario valence and affective impact (see Supplemental Methods and Table S1). The experimental paradigm and data acquisition across both experiments were controlled via PsychoPy (Version 2020.2.4).

### MRI Data Acquisition and Preprocessing

Neuroimaging data were acquired using a 3T Siemens Prisma scanner (Siemens Healthineers, Erlangen, Germany) equipped with a 32-channel head coil at the University of Macau. Data preprocessing was conducted using SPM12 (Wellcome Department of Imaging Neuroscience, London, UK) within the MATLAB R2021b environment. More details were provided in the Supplementary Methods.

### Behavioral Data Analysis

Behavioral data from both experiments were analyzed on future likelihood prediction using LMMs implemented via the *lme4* package (51) in R (Version 4.4.0). Across both experiments, the models included Agent, Valence, and SA as fixed effects, along with their fully crossed interaction terms. In Experiment 1, SA was entered as a categorical variable (high versus low), whereas in Experiment 2, SA was entered as a continuous variable using the total LSAS score. To account for the hierarchical structure of the repeated-measures data and to control for individual differences in baseline ratings, a random intercept for each participant was included in all models. To maintain conservative and reliable statistical inferences, degrees of freedom were calculated using the Kenward-Roger approximation. Data visualization was conducted using *ggplot2* package (50) in R.

### fMRI Data Analysis

#### MVPA Decoding

To capture fine-grained neural patterns, we employed the Least Squares All (LSA) approach in SPM12 (51). For each participant, a GLM was constructed where each individual trial was modeled as a separate regressor convolved with the canonical Hemodynamic Response Function (HRF). Six head-motion parameters were included as nuisance regressors. This process yielded a series of single-trial beta maps representing the spatial activation patterns for Self and Other conditions.

The ROIs of social cognitive network was guided by a meta-analysis of mentalizing studies (21). Using the MarsBaR toolbox (52), we created six spherical ROIs with a 10-mm radius centered the following MNI coordinates (Figure S1): the mPFC [1, 58, 19], precuneus [3, –56, 37], left TPJ [–52, –56, 24], right TPJ [55, –53, 24], left STS [–59, –26, –9], and right STS [59, –18, –17]. The bilateral pgACC and bilateral AI (Figure S1) were extracted with their anatomical masks defined using the AAL3 atlas (53). Using these masks, we extracted voxel-wise spatial patterns from the unsmoothed single-trial beta maps. The extracted data were organized into a multi-dimensional dataset to serve as the input for the subsequent multivariate decoding analysis.

Decoding was performed using the CoSMoMVPA toolbox (54) implemented in MATLAB. A Linear Discriminant Analysis (LDA) classifier was trained to distinguish between the neural patterns of the target pairs, such as self-negative versus other-negative trials (see Supplemental Methods). Group-level decoding accuracy was tested against the theoretical chance level (0.5) using a one-sample *t*-test (one-tailed). *P*-values were then corrected for multiple comparisons across all ROIs using the False Discovery Rate (FDR) method.

#### Parametric Modulation Analysis

Individual-level GLMs modeled the four primary experimental conditions (self-positive, self-negative, other-positive, and other-negative; refer to Supplementary Methods for detail). To identify brain regions dynamically tracking subjective evaluations, trial-by-trial ratings of social acceptance, perceived anxiety, and future likelihood prediction were included as mean-centered parametric modulators. Crucially, we utilized serial orthogonalization, entering likelihood prediction last. This hierarchical approach isolated neural activity specifically associated with likelihood prediction, controlling for general affective factors (i.e., perceived acceptance and anxiety).

At the group level, one sample *t*-tests were conducted to examine the main effects of each condition and their respective parametric modulators (i.e., social acceptance, perceived anxiety, and likelihood estimation). Subsequently, we performed paired *t*-tests to directly compare the neural sensitivity between self- and other-referential processing (e.g., self-negative versus other-negative predictions). Group results were thresholded at *p*_uncorrected_ < .001 with cluster-level family-wise error (FWE) correction *p*_FWE_ < .05 to control for multiple comparisons.

### Brain-Behavior Integration Analysis

To evaluate the contribution of neural representational distinctions to SA severity (LSAS total score), we performed multiple linear regression models using ROI decoding accuracies as predictors separately for positive and negative conditions. Next, to link neural representation with functional sensitivity to social predictions, we conducted FDR-corrected Spearman correlation analyses between the ROI decoding accuracies (MVPA) and the parametric modulation beta weights (GLM).

Guided by these correlations, we conducted a mediation analysis to investigate whether the representational distinctiveness in the left pgACC explained the relationship between SA severity and the precuneus’s computational sensitivity to self-versus other-negative predictions. The mediation model evaluated the indirect effect using non-parametric bootstrapping (5,000 iterations). Detailed model specifications are provided in the Supplementary Methods.

## Acknowledgments

This research received funding from National Natural Science Foundation of China (32000733), Humanities and Social Sciences Research Project of the Ministry of Education (24YJA190011), Henan Province Key Scientific Research Project Plan for Higher Education Institutions (25A190002). The funders had no role in the design, data collection, data analysis, and reporting of this study. The data and code that support the findings of this study are available at https://osf.io/xt9r7/overview?view_only=f7dfe35654864e4cbffcc3db5e379a52. The authors acknowledge the use of Gemini (Google LLC) for English language proofreading and code debugging during the preparation of this work.

## Author contributions

**Yanyan Qi:** Conceptualization, Methodology, Investigation, Data Curation, formal analysis, Validation, Visualization, Writing - Original Draft, Writing - Review & Editing; Funding acquisition. **Yun Chen:** Methodology, Software, Writing - Review & Editing. **Yixiao Hu:** Visualization, Writing - Review & Editing; **Qianqian Fan:** Investigation, Writing - Review & Editing. **Lingxiao Wang:** Methodology, Formal analysis. **Guochun Yang:** Formal analysis, Validation, Visualization, Writing - Review & Editing. **Grit Hein:** Conceptualization, Validation, Writing - Review & Editing**; Haiyan Wu:** Project administration, Resources, Funding acquisition, Conceptualization, Methodology, Validation, Writing – review & editing, Supervision.

## Declaration of interest

The authors declare no competing interests.

## Notes

### Competing Interest Statement

The authors have declared no competing interest.

### Summary of Updates

Supplemental files updated. Funding information updated.

## References

1. Beer JS. Exaggerated Positivity in Self-Evaluation: A Social Neuroscience Approach to Reconciling the Role of Self-esteem Protection and Cognitive Bias. Social and Personality Psychology Compass. 2014;8(10):583–94. doi:10.1111/spc3.12133

2. Sharot T. The optimism bias. Current Biology. 2011 Dec 6;21(23):R941–5. doi:10.1016/j.cub.2011.10.030

3. Weinstein ND. Unrealistic optimism about future life events. Journal of Personality and Social Psychology. 1980;39(5):806–20. doi:10.1037/0022-3514.39.5.806

4. Blair KS, Otero M, Teng C, Geraci M, Ernst M, Blair RJR, et al. Reduced optimism and a heightened neural response to everyday worries are specific to generalized anxiety disorder, and not seen in social anxiety. Psychological Medicine. 2017 Jul;47(10):1806–15. doi:10.1017/S0033291717000265

5. Karnick AT, Brick LA. From perception to projection: Exploring neuroaffective advances in understanding optimism bias and belief updating. Current Opinion in Psychology. 2024 Dec 1;60:101937. doi:10.1016/j.copsyc.2024.101937

6. Basanovic J, Kowal L, Millward S, MacLeod C. The legacy of social anxiety-linked negative expectancy: A pathway from pre-event negative expectancies to post-event negative thinking. Journal of Behavior Therapy and Experimental Psychiatry. 2024 Jun 1;83:101937. doi:10.1016/j.jbtep.2023.101937

7. Cao J, Gu R, Bi X, Zhu X, Wu H. Unexpected Acceptance? Patients with Social Anxiety Disorder Manifest their Social Expectancy in ERPs During Social Feedback Processing. Front Psychol. 2015 Nov 16;6. doi:10.3389/fpsyg.2015.01745

8. Qi Y, Gu R, Cao J, Bi X, Wu H, Liu X. Response bias-related impairment of early subjective face discrimination in social anxiety disorders: An event-related potential study. Journal of Anxiety Disorders. 2017 Apr 1;47:10–20. doi:10.1016/j.janxdis.2017.02.003

9. Blair K, Geraci M, Devido J, McCaffrey D, Chen G, Vythilingam M, et al. Neural Response to Self- and Other Referential Praise and Criticism in Generalized Social Phobia. Archives of General Psychiatry. 2008 Oct 6;65(10):1176–84. doi:10.1001/archpsyc.65.10.1176

10. Blair KS, Otero M, Teng C, Jacobs M, Odenheimer S, Pine DS, et al. Dissociable roles of ventromedial prefrontal cortex (vmPFC) and rostral anterior cingulate cortex (rACC) in value representation and optimistic bias. NeuroImage. 2013 Sep 1;78:103–10. doi:10.1016/j.neuroimage.2013.03.063

11. Fields EC, Weber K, Stillerman B, Delaney-Busch N, Kuperberg GR. Functional MRI reveals evidence of a self-positivity bias in the medial prefrontal cortex during the comprehension of social vignettes. Soc Cogn Affect Neurosci. 2019 Aug 7;14(6):613–21. doi:10.1093/scan/nsz035

12. Finlayson-Short L, Harrison BJ, Davey C. Self-other referential neural processing in social anxiety disorder and major depressive disorder. NeuroImage: Clinical. 2021 Jan 1;30:102669. doi:10.1016/j.nicl.2021.102669

13. Hughes BL, Beer JS. Protecting the self: the effect of social-evaluative threat on neural representations of self. J Cogn Neurosci. 2013 Apr;25(4):613–22. doi:10.1162/jocn_a_00343 PubMed PMID: 23249346.

14. Sharot T, Riccardi AM, Raio CM, Phelps EA. Neural mechanisms mediating optimism bias. Nature. 2007 Nov;450(7166):102–5. doi:10.1038/nature06280

15. Sharot T, Korn CW, Dolan RJ. How unrealistic optimism is maintained in the face of reality. Nat Neurosci. 2011 Nov;14(11):1475–9. doi:10.1038/nn.2949

16. Sharot T, Rollwage M, Sunstein CR, Fleming SM. Why and When Beliefs Change. Perspect Psychol Sci. 2023 Jan;18(1):142–51. doi:10.1177/17456916221082967 PubMed PMID: 35939828.

17. Menon V. 20 years of the default mode network: A review and synthesis. Neuron. 2023 Aug 16;111(16):2469–87. doi:10.1016/j.neuron.2023.04.023 PubMed PMID: 37167968.

18. Northoff G, Heinzel A, de Greck M, Bermpohl F, Dobrowolny H, Panksepp J. Self-referential processing in our brain--a meta-analysis of imaging studies on the self. Neuroimage. 2006 May 15;31(1):440–57. doi:10.1016/j.neuroimage.2005.12.002 PubMed PMID: 16466680.

19. Qin P, Northoff G. How is our self related to midline regions and the default-mode network? NeuroImage. 2011 Aug 1;Special Issue: Educational Neuroscience57(3):1221–33. doi:10.1016/j.neuroimage.2011.05.028

20. Schurz M, Radua J, Aichhorn M, Richlan F, Perner J. Fractionating theory of mind: A meta-analysis of functional brain imaging studies. Neuroscience & Biobehavioral Reviews. 2014 May 1;42:9–34. doi:10.1016/j.neubiorev.2014.01.009

21. van Veluw SJ, Chance SA. Differentiating between self and others: an ALE meta-analysis of fMRI studies of self-recognition and theory of mind. Brain Imaging and Behavior. 2014 Mar 1;8(1):24–38. doi:10.1007/s11682-013-9266-8

22. Courtney AL, Meyer ML. Self-Other Representation in the Social Brain Reflects Social Connection. J Neurosci. 2020 Jul 15;40(29):5616–27. doi:10.1523/JNEUROSCI.2826-19.2020 PubMed PMID: 32541067.

23. Parkinson C, Kleinbaum AM, Wheatley T. Similar neural responses predict friendship. Nat Commun. 2018 Jan 30;9(1):332. doi:10.1038/s41467-017-02722-7

24. Shen YL, Hyon R, Wheatley T, Kleinbaum AM, Welker CL, Parkinson C. Neural similarity predicts whether strangers become friends. Nat Hum Behav. 2025 Nov;9(11):2285–98. doi:10.1038/s41562-025-02266-7

25. Mitchell JP, Macrae CN, Banaji MR. Dissociable Medial Prefrontal Contributions to Judgments of Similar and Dissimilar Others. Neuron. 2006 May 18;50(4):655–63. doi:10.1016/j.neuron.2006.03.040 PubMed PMID: 16701214.

26. Wang Y, Wang R, Wu H. The role of oxytocin in modulating self–other distinction in human brain: a pharmacological fMRI study. Cerebral Cortex. 2023 Mar 1;33(5):1708–25. doi:10.1093/cercor/bhac167

27. Wu H, Yang S, Sun S, Liu C, Luo Y jia. The Male Advantage in Child Facial Resemblance Detection: Behavioral and ERP evidence. Soc Neurosci. 2013 Nov;8(6):. doi:10.1080/17470919.2013.835279 PubMed PMID: 24053135; PubMed Central PMCID: PMC3815694.

28. Boehme S, Miltner WHR, Straube T. Neural correlates of self-focused attention in social anxiety. Soc Cogn Affect Neurosci. 2015 Jun 1;10(6):856–62. doi:10.1093/scan/nsu128

29. Leichsenring F, Leweke F. Social Anxiety Disorder. New England Journal of Medicine. 2017 Jun 8;376(23):2255–64. doi:10.1056/NEJMcp1614701 PubMed PMID: 28591542.

30. Morrison AS, Heimberg RG. Social Anxiety and Social Anxiety Disorder. Annual Review of Clinical Psychology. 2013 Mar 28;9(Volume 9, 2013):249–74. doi:10.1146/annurev-clinpsy-050212-185631

31. Spurr JM, Stopa L. Self-focused attention in social phobia and social anxiety. Clinical Psychology Review. 2002 Sep 1;22(7):947–75. doi:10.1016/S0272-7358(02)00107-1

32. Yoon HJ, Seo EH, Kim JJ, Choo IH. Neural Correlates of Self-referential Processing and Their Clinical Implications in Social Anxiety Disorder. Clin Psychopharmacol Neurosci. 2019 Feb;17(1):12–24. doi:10.9758/cpn.2019.17.1.12 PubMed PMID: 30690936; PubMed Central PMCID: PMC6361035.

33. Decety J, Sommerville J. Shared Representations between Self and Other: A Social Cognitive Neuroscience View. Trends in cognitive sciences. 2004 Jan 1;7:527–33. doi:10.1016/j.tics.2003.10.004

34. Hein G, Qi Y, Han S. The biological foundations and modulation of empathy. In: Social psychology: Handbook of basic principles, 3rd ed. New York, NY, US: The Guilford Press; 2021. p. 127–39.

35. Lamm C, Bukowski H, Silani G. From shared to distinct self–other representations in empathy: evidence from neurotypical function and socio-cognitive disorders. Philos Trans R Soc Lond B Biol Sci. 2016 Jan 19;371(1686):20150083. doi:10.1098/rstb.2015.0083

36. Gao W, Li Y, Yuan J, He Q. The Shared and Distinct Mechanisms Underlying Fear of Evaluation in Social Anxiety: The Roles of Negative and Positive Evaluation. Depression and Anxiety. 2025;2025(1):9559056. doi:10.1155/da/9559056

37. Weeks JW, Heimberg RG, Rodebaugh TL, Norton PJ. Exploring the relationship between fear of positive evaluation and social anxiety. Journal of Anxiety Disorders. 2008 Apr 1;22(3):386–400. doi:10.1016/j.janxdis.2007.04.009

38. Guterstam A, Bio BJ, Wilterson AI, Graziano M. Temporo-parietal cortex involved in modeling one’s own and others’ attention. Peelen MV, Frank MJ, Atkinson A, editors. eLife. 2021 Feb 15;10:e63551. doi:10.7554/eLife.63551

39. Apps MAJ, Rushworth MFS, Chang SWC. The Anterior Cingulate Gyrus and Social Cognition: Tracking the Motivation of Others. Neuron. 2016 May 18;90(4):692–707. doi:10.1016/j.neuron.2016.04.018 PubMed PMID: 27196973.

40. Eisenberger NI, Lieberman MD, Williams KD. Does Rejection Hurt? An fMRI Study of Social Exclusion. Science. 2003 Oct 10;302(5643):290–2. doi:10.1126/science.1089134

41. Hein G, Silani G, Preuschoff K, Batson CD, Singer T. Neural Responses to Ingroup and Outgroup Members’ Suffering Predict Individual Differences in Costly Helping. Neuron. 2010 Oct 6;68(1):149–60. doi:10.1016/j.neuron.2010.09.003 PubMed PMID: 20920798.

42. Somerville LH, Heatherton TF, Kelley WM. Anterior cingulate cortex responds differentially to expectancy violation and social rejection. Nat Neurosci. 2006 Aug;9(8):1007–8. doi:10.1038/nn1728

43. Schade E, White PJ, Maramaldi P. Self-consciousness in adolescent and adult social anxiety: A systematic review of fMRI and EEG findings. Biological Psychology. 2025 Nov 1;202:109137. doi:10.1016/j.biopsycho.2025.109137

44. Mizzi S, Pedersen M, Rossell SL, Rendell P, Terrett G, Heinrichs M, et al. Resting-state amygdala subregion and precuneus connectivity provide evidence for a dimensional approach to studying social anxiety disorder. Transl Psychiatry. 2024 Mar 14;14(1):1–10. doi:10.1038/s41398-024-02844-9

45. Korn CW, Prehn K, Park SQ, Walter H, Heekeren HR. Positively Biased Processing of Self-Relevant Social Feedback. J Neurosci. 2012 Nov 21;32(47):16832–44. doi:10.1523/JNEUROSCI.3016-12.2012 PubMed PMID: 23175836.

46. Cavanna AE, Trimble MR. The precuneus: a review of its functional anatomy and behavioural correlates. Brain. 2006 Mar 1;129(3):564–83. doi:10.1093/brain/awl004

47. Dadario NB, Sughrue ME. The functional role of the precuneus. Brain. 2023 Sep 1;146(9):3598–607. doi:10.1093/brain/awad181

48. Sohail A, Zhang L. Informing the treatment of social anxiety disorder with computational and neuroimaging data. psychoradiology. 2024 Mar 1;4:kkae010. doi:10.1093/psyrad/kkae010

49. Liebowitz MR. Social phobia. Modern problems of pharmacopsychiatry. 1987;22(141):e173.

50. Wickham H. ggplot2. Wiley interdisciplinary reviews: computational statistics. 2011;3(2):180–5.

51. Rissman J, Gazzaley A, D’Esposito M. Measuring functional connectivity during distinct stages of a cognitive task. Neuroimage. 2004;23(2):752–63.

52. Brett M, Anton JL, Valabregue R, Poline JB. Region of interest analysis using an SPM toolbox. In: Vol. 16. Sendai; 2002. p. 497.

53. Rolls ET, Huang CC, Lin CP, Feng J, Joliot M. Automated anatomical labelling atlas 3. Neuroimage. 2020;206:116189.

54. Oosterhof NN, Connolly AC, Haxby JV. CoSMoMVPA: multi-modal multivariate pattern analysis of neuroimaging data in Matlab/GNU Octave. Frontiers in neuroinformatics. 2016;10:27.

