## Supplemental files for "Self-Other Neural Representational Distinction is Associated with Distorted Social Predictions in Social Anxiety"

**Supplementary materials**

**Supplementary Figure**

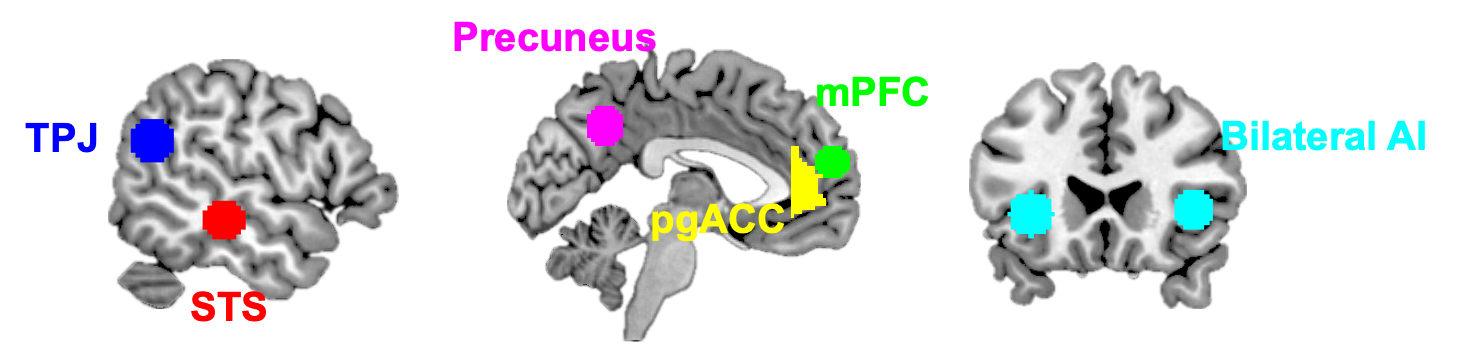

**Figure S1. Regions of interest (ROIs) for multivariate pattern analysis decoding.** Ten ROIs were selected based on *a priori* hypotheses involving the Theory of Mind (ToM) network and social-affective processing. Six spherical ROIs (10-mm radius) were centered on coordinates derived from a meta-analysis of mentalizing studies (van Veluw & Chance, 2014), including the medial prefrontal cortex (mPFC), precuneus, bilateral temporoparietal junction (TPJ), and bilateral superior temporal sulcus (STS). To capture neural patterns associated with social evaluation, anatomical masks for the bilateral pregenual anterior cingulate cortex (pgACC) and bilateral anterior insula (AI) were defined using the AAL3 atlas (Rolls et al., 2020). Here the TPJ, STS and pgACC on the left side were shown.

**Supplementary Tables**

**Table S1**. Descriptive statistics for the psychological features of positive and negative social scenarios

| **Condition** | **Set** | **N** | **Arousal (M±SD)** | **Valence (M±SD)** | **Pleasantness (M±SD)** | **Acceptance (M±SD)** |
| --- | --- | --- | --- | --- | --- | --- |
| **Positive** | Full Set (27 items) | 27 | 6.733 ± 0.351 | 2.031 ± 0.485 | 2.782 ± 0.231 | 2.831 ± 0.291 |
|  | Subset  (20 items) | 20 | 5.652 ± 0.147 | 2.439 ± 0.269 | 2.778 ± 0.265 | 2.810 ± 0.335 |
| **Negative** | Full Set (27 items) | 27 | 5.777 ± 0.634 | -2.094 ± 0.574 | -2.282 ± 0.462 | -2.094 ± 0.578 |
|  | Subset  (20 items) | 20 | 4.993 ± 0.319 | -1.967 ± 0.447 | -2.141 ± 0.477 | -2.028 ± 0.471 |

*Note*: *M* represents the mean; *SD* represents the standard deviation. Arousal was rated on a 1–9 scale, while Valence, Pleasantness, and Acceptance were rated on a −4 to 4 scale. The initial 4 participants completed the task using the “Full Set” (27 items), while the subsequent 32 participants completed the task using the “Subset” (20 items), which consisted of the first 20 items from each valence category of the full set.

**Table S2.** Linear Mixed-Effects Model for Future Likelihood Prediction in Experiment 1

|  | ***χ^2^* (ANOVA-Type III)** | ***df*** | ***p*** | ***b*** | ***SE*** |
| --- | --- | --- | --- | --- | --- |
| **A LMM testing the three-way interaction effect of Agent x Valence x Social Anxiety (high vs. low)** | | | | | |
| (Intercept) | 3314.735 | 1 | <0.001 | 53.752 | 0.934 |
| Agent | 10.295 | 1 | <0.001 | 1.213 | 0.378 |
| Valence | 4053.091 | 1 | <0.001 | -24.058 | 0.378 |
| Social Anxiety | 0.013 | 1 | 0.910 | 0.210 | 1.876 |
| Agent x Valence | 60.980 | 1 | <0.001 | -5.902 | 0.756 |
| Agent x Social Anxiety | 0.091 | 1 | 0.763 | 0.228 | 0.756 |
| Valence x Social Anxiety | 253.597 | 1 | <0.001 | 12.036 | 0.756 |
| Agent x Valence x Social Anxiety | 102.724 | 1 | <0.001 | -15.320 | 1.512 |
| **B LMM testing the two-way interaction effect of Agent x Social Anxiety (high vs. low) in positive condition** | | | | | |
| (Intercept) | 3660.531 | 1 | <0.001 | 65.781 | 1.087 |
| Agent | 75.167 | 1 | <0.001 | 4.163 | 0.480 |
| Social Anxiety | 7.133 | 1 | 0.008 | -5.808 | 2.175 |
| Agent x Social Anxiety | 67.456 | 1 | <0.001 | 7.888 | 0.960 |
| **C LMM testing the twp-way interaction effect of Agent x Social Anxiety (high vs. low) in negative condition** | | | | | |
| (Intercept) | 1049.325 | 1 | <0.001 | 41.723 | 1.288 |
| Agent | 11.421 | 1 | <0.001 | -1.739 | 0.514 |
| Social Anxiety | 5.845 | 1 | 0.016 | 6.228 | 2.576 |
| Agent x Social Anxiety | 52.183 | 1 | <0.001 | -7.432 | 1.029 |

**Table S3.** Linear Mixed-Effects Model for Future Likelihood Prediction in Experiment 2

|  | ***χ^2^* (ANOVA-Type III)** | ***df*** | ***p*** | ***b*** | ***SE*** |
| --- | --- | --- | --- | --- | --- |
| **A LMM testing the three-way interaction effect of Agent x Valence x Social Anxiety (continuous variable)** | | | | | |
| (Intercept) | 112.534 | 1 | <0.001 | 40.115 | 3.782 |
| Agent | 16.431 | 1 | <0.001 | -8.422 | 2.078 |
| Valence | 239.047 | 1 | <0.001 | 31.934 | 2.065 |
| Social Anxiety | 0.475 | 1 | 0.491 | 0.057 | 0.082 |
| Agent x Valence | 13.651 | 1 | <0.001 | 10.808 | 2.925 |
| Agent x Social Anxiety | 3.281 | 1 | 0.07 | 0.082 | 0.045 |
| Valence x Social Anxiety | 11.833 | 1 | <0.001 | -0.155 | 0.045 |
| Agent x Valence x Social Anxiety | 4.198 | 1 | 0.04 | -0.131 | 0.064 |
| **B LMM testing the two-way interaction effect of Agent x Social Anxiety in positive condition** | | | | | |
| (Intercept) | 167.678 | 1 | <0.001 | 71.824 | 5.547 |
| Agent | 1.834 | 1 | 0.176 | 2.403 | 1.774 |
| Social Anxiety | 0.623 | 1 | 0.430 | -0.095 | 0.120 |
| Agent x Social Anxiety | 1.645 | 1 | 0.200 | -0.050 | 0.039 |
| **C LMM testing the twp-way interaction effect of Agent x Social Anxiety in negative condition** | | | | | |
| (Intercept) | 74.351 | 1 | <0.001 | 40.306 | 4.674 |
| Agent | 16.502 | 1 | <0.001 | -8.366 | 2.059 |
| Social Anxiety | 0.261 | 1 | 0.610 | 0.052 | 0.102 |
| Agent x Social Anxiety | 3.417 | 1 | 0.065 | 0.083 | 0.045 |

*Note.* Social Anxiety, LSAS total score**,** df = degrees of freedom for Wald Chi-square tests. For regression coefficients *b*, the reference levels were “Other” and “Negative”. SE = standard error. Statistical significance was determined using Type III Wald chi-square tests.

**Table S4.** Decoding accuracy of self versus other in positive and negative social contexts

|  | **Mean Accuracy** | ***SD*** | ***95% CI.low*** | ***p_uncorrected_*** | ***p_FDR_*** |
| --- | --- | --- | --- | --- | --- |
| **Positive: Self versus Other** |  |  |  |  |  |
| lAI | 0.536 | 0.111 | 0.504 | 0.031 | 0.052 |
| l-pgACC | 0.538 | 0.112 | 0.507 | 0.025 | 0.051 |
| lSTS | 0.528 | 0.120 | 0.494 | 0.088 | 0.098 |
| lTPJ | 0.540 | 0.122 | 0.505 | 0.031 | 0.052 |
| mPFC | 0.538 | 0.125 | 0.502 | 0.041 | 0.063 |
| precuneus | 0.547 | 0.120 | 0.513 | 0.013 | 0.051 |
| rAI | 0.523 | 0.113 | 0.490 | 0.119 | 0.125 |
| r-pgACC | 0.537 | 0.104 | 0.508 | 0.021 | 0.051 |
| rSTS | 0.525 | 0.133 | 0.487 | 0.135 | 0.135 |
| rTPJ | 0.553 | 0.108 | 0.522 | 0.003 | 0.034 |
| **Negative: Self versus Other** |  |  |  |  |  |
| lAI | 0.544 | 0.130 | 0.507 | 0.026 | 0.051 |
| l-pgACC | 0.526 | 0.111 | 0.495 | 0.084 | 0.098 |
| lSTS | 0.565 | 0.118 | 0.532 | 0.0012 | 0.025 |
| lTPJ | 0.534 | 0.125 | 0.498 | 0.058 | 0.083 |
| mPFC | 0.528 | 0.106 | 0.498 | 0.063 | 0.084 |
| precuneus | 0.530 | 0.118 | 0.500 | 0.069 | 0.086 |
| rAI | 0.541 | 0.107 | 0.511 | 0.014 | 0.051 |
| r-pgACC | 0.541 | 0.120 | 0.510 | 0.017 | 0.051 |
| rSTS | 0.534 | 0.096 | 0.506 | 0.022 | 0.051 |
| rTPJ | 0.548 | 0.117 | 0.514 | 0.011 | 0.051 |

*Note*. l-/ r- pgACC: left/ right pregenual anterior cingulate cortex; lAI/ rAI: left/ right anterior insula; lSTS/ rSTS: left/ right superior temporal sulcus; lTPJ/ rTPJ: left/ right temporoparietal junction; mPFC: medial prefrontal cortex. 95% CI.low: the low bound of 95% confidence interval (CI). We only presented the lower bound of CI as a one-tailed *t*-test was employed. *p* values were corrected using False Discovery Rate (FDR) method across all ROIs and all conditions (i.e., Positive and Negative valence).

**Table S5.** A linear regression testing the association between decoding accuracy and social anxiety (LSAS total score) in positive condition

|  | ***b*** | ***SE*** | ***T*** | ***p*** |
| --- | --- | --- | --- | --- |
| (Intercept) | 46.455 | 27.373 | 1.697 | 0.104 |
| l-pgACC | 74.364 | 34.634 | 2.147 | 0.043 |
| r-pgACC | 7.122 | 31.552 | 0.226 | 0.824 |
| lTPJ | -2.918 | 39.085 | -0.075 | 0.941 |
| lAI | 11.474 | 25.825 | 0.444 | 0.661 |
| lSTS | -33.128 | 35.97 | -0.921 | 0.367 |
| mPFC | -16.586 | 27.411 | -0.605 | 0.551 |
| precuneus | -58.256 | 33.557 | -1.736 | 0.097 |
| rAI | 9.486 | 32.376 | 0.293 | 0.772 |
| rSTS | -37.662 | 26.576 | -1.417 | 0.170 |
| rTPJ | 39.668 | 44.837 | 0.885 | 0.386 |

*Note*. l-/ r- pgACC: left/ right pregenual anterior cingulate cortex; lAI/ rAI: left/ right anterior insula; lSTS/ rSTS: left/ right superior temporal sulcus; lTPJ/ rTPJ: left/ right temporoparietal junction; mPFC: medial prefrontal cortex.

**Table S6.** A linear regression testing the association between decoding accuracy and social anxiety (LSAS total score) in negative condition

|  | ***b*** | ***SE*** | ***T*** | ***p*** |
| --- | --- | --- | --- | --- |
| (Intercept) | 42.244 | 22.349 | 1.890 | 0.072 |
| l-pgACC | 93.024 | 34.059 | 2.731 | 0.012 |
| r-pgACC | -13.052 | 39.109 | -0.334 | 0.742 |
| lTPJ | -59.976 | 29.134 | -2.059 | 0.052 |
| lAI | -20.236 | 25.103 | -0.806 | 0.429 |
| lSTS | 10.536 | 31.738 | 0.332 | 0.743 |
| mPFC | -10.082 | 36.059 | -0.280 | 0.782 |
| precuneus | 44.48 | 28.398 | 1.566 | 0.132 |
| rAI | -44.943 | 38.235 | -1.175 | 0.252 |
| rSTS | 34.766 | 32.237 | 1.079 | 0.293 |
| rTPJ | -28.988 | 37.290 | -0.777 | 0.445 |

*Note*. l-/ r- pgACC: left/ right pregenual anterior cingulate cortex; lAI/ rAI: left/ right anterior insula; lSTS/ rSTS: left/ right superior temporal sulcus; lTPJ/ rTPJ: left/ right temporoparietal junction; mPFC: medial prefrontal cortex.

**Table S7**. Brain regions showing significant activation for the contrast of Self versus Other in positive context

| **Anatomical Region** | **Hemi** | **Peak MNI**  **(x, y, z)** | **Cluster Size**  **(voxel)** | **Peak T** | ***p*_FWE_ (cluster)** |
| --- | --- | --- | --- | --- | --- |
| Medial Prefrontal Cortex (mPFC) | R | 12, 54, 14 | 631 | 5.31 | < 0.001 |
| Anterior Cingulate Cortex (ACC) | L | -6, 10, 22 | 233 | 5.42 | 0.001 |
| Caudate / Striatum | R | 14, 0, 22 | 229 | 4.84 | 0.001 |

**Table S8.** Univariate regression: Social anxiety on neural self-other activation differences in positive context

| **Anatomical Region** | **Hemi** | **MNI (x, y, z)** | **Cluster Size (voxel)** | **Peak *T*** | ***p*_uncorrected_** |
| --- | --- | --- | --- | --- | --- |
| Middle Frontal Gyrus | R | 34, 26, 38 | 75 | 4.49 | < 0.001 |
| Pregenual Anterior Cingulate Cortex | L | -12, 34, 14 | 9 | 4.11 | < 0.001 |
| Superior Frontal Gyrus | R | 36, 18, 58 | 44 | 3.79 | < 0.001 |
| Medial Frontal Gyrus | R | 10, 30, 34 | 15 | 3.57 | 0.001 |
| Insula / Postcentral | L | -30, -34, 26 | 23 | 3.56 | 0.001 |
| Middle Temporal Gyrus | R | 32, -62, 0 | 13 | 3.49 | 0.001 |
| Inferior Frontal Gyrus | R | 50, 40, -2 | 65 | 3.42 | 0.001 |
| Inferior Frontal Gyrus | L | -20, 36, -6 | 17 | 3.41 | 0.001 |
| Middle Temporal Gyrus | L | -32, -60, 10 | 13 | 3.33 | 0.001 |
| Postcentral Gyrus | R | 22, -32, 28 | 12 | 3.28 | 0.001 |
| Right Angular Gyrus | R | 36, -48, 30 | 21 | 3.27 | 0.001 |
| Subgenual Anterior Cingulate Cortex | L | -12, 26, -4 | 15 | 3.24 | 0.001 |
| Middle Frontal Gyrus | L | -30, 26, 56 | 47 | 3.23 | 0.001 |
| Middle Frontal Gyrus | L | -36, 40, 16 | 9 | 3.11 | 0.002 |
| Posterior Cingulate Cortex | R | 16, -40, 22 | 9 | 3.01 | 0.003 |

**Table S9.** Univariate regression: Social anxiety on neural self-other activation differences (negative outcomes).

| **Anatomical Region** | **Hemi** | **MNI (x, y, z)** | **Cluster Size (voxel)** | **Peak *T*** | ***p*_uncorrected_** |
| --- | --- | --- | --- | --- | --- |
| Middle Frontal Gyrus | L | -32, 36, 16 | 10 | 3.73 | < .001 |
| Insula | R | 38, -34, -2 | 14 | 3.37 | 0.001 |
| Middle Frontal Gyrus | L | -28, 42, 8 | 32 | 3.29 | 0.001 |

**Table S10. Spearman correlations between regional decoding accuracies (self versus other) and precuneus neural tracking of negative social predictions.**

|  | Precuneus Neural Tracking  (Self versus Other Negative) | | |
| --- | --- | --- | --- |
|  | *r* | *p_uncorrected_* | *p_FDR_* |
| Positive: Self versus Other Decoding Accuracy | | | |
| l-pgACC | 0.627 | < 0.001 | 0.003 |
| r-pgACC | 0.120 | 0.521 | 0.694 |
| ITPJ | 0.140 | 0.452 | 0.646 |
| IAI | 0.161 | 0.387 | 0.645 |
| ISTS | 0.195 | 0.294 | 0.598 |
| mPFC | 0.446 | 0.012 | 0.079 |
| precuneus | 0.330 | 0.070 | 0.350 |
| rAI | 0.107 | 0.565 | 0.707 |
| rSTS | 0.199 | 0.284 | 0.598 |
| rTPJ | 0.453 | 0.010 | 0.079 |
| Negative: Self versus other Decoding Accuracy | | | |
| l-pgACC | 0.221 | 0.233 | 0.598 |
| r-pgACC | -0.195 | 0.294 | 0.598 |
| ITPJ | -0.032 | 0.864 | 0.864 |
| IAI | -0.173 | 0.351 | 0.638 |
| ISTS | 0.071 | 0.703 | 0.812 |
| mPFC | -0.062 | 0.739 | 0.812 |
| precuneus | 0.280 | 0.127 | 0.507 |
| rAI | -0.146 | 0.434 | 0.646 |
| rSTS | -0.193 | 0.299 | 0.598 |
| rTPJ | 0.054 | 0.771 | 0.812 |

*Note*: l- /r- pgACC: left/ right pregenual anterior cingulate cortex; lTPJ/ rTPJ: left/ right temporoparietal junction; lAI/ rAI: left/ right anterior insula; lSTS/ rSTS: left/ right superior temporal sulcus; mPFC: medial prefrontal cortex. *p* values were corrected using False Discovery Rate (FDR) method across all ROIs and all conditions (i.e., Positive and Negative valence).

**Supplementary Methods**

*A Priori Power Analyses for Sample Size Determination*

For Experiment 1, an a priori power analysis was performed using G*Power (Version 3.1.9.7) for a repeated-measures ANOVA with a within-between interaction design (two within-subject factors: Agent and Valence; one between-subject factor: SA). Assuming α = 0.05, a statistical power of 95%, and a medium effect size (Cohen’s *f*^2^ = 0.20), a minimum of 28 participants per group was required.

For Experiment 2, a similar a priori power analysis indicated that a sample size of 34 participants would be sufficient to detect a medium effect size with 80% power at α = 0.05 for a repeated-measures ANOVA (two within-subject factors: Agent and Valence). This calculation specifically targeted the factorial main effects and interactions.

*Experimental Design and Experimental Materials*

The experimental stimuli consisted of a validated set of social interaction scenarios. Initially, a broader pool of situations was evaluated by an independent panel (21 females, N = 41, Mage = 21.073, SD = 1.673 ) across multiple affective dimensions, including valence, pleasantness, and emotional arousal. From this pool, 54 scenarios were selected for the final task, balanced for valence: 27 depicted favorable social outcomes (e.g., “Receiving a warm ovation after a public speech”), while 27 described adverse social experiences (e.g., “Facing a lack of support during academic challenges”).

In Experiment 1, all participants completed the full set of 54 scenarios under both the Self and Other conditions, yielding a total of 108 trials. In Experiment 2, to ensure high data quality and minimize participant fatigue during the fMRI scanning session, the task length was streamlined following the initial recruitment phase. The first four participants completed the original 108-trial version (6 blocks). For the remaining 32 participants, the task was optimized to 80 trials (4 blocks; 20 trials per condition). This optimized version utilized a representative subset of 40 scenarios (20 positive, 20 negative) selected from the original set to maintain consistent affective impact and psychometric properties (see Supplementary Table S1). Critically, the task logic, trial-level timing, and experimental balance remained strictly identical across all participants. To account for the variation in trial numbers during group-level statistical analysis, all behavioral and neural metrics were either averaged or modeled as proportion-based weights to ensure equal contribution from each participant.

Descriptive statistics for the emotional features (Arousal, Valence, Pleasantness, and Acceptance) of both the original 27-item set and the final 20-item set are provided below (Table S1). These ratings confirm that the shortened set maintained the intended emotional properties of the original materials.

**Positive social scenarios**

1. After presenting the report, “I/Liu Yang received applause.”
2. After participating in the competition, “I/Liu Yang received applause.”
3. When the teacher required group projects, “I/Liu Yang received many invitations from my classmates.”
4. I/Liu Yang went to a strange place to collect materials and “was warmly received”.
5. “Everyone around me/Liu Yang likes me very much.”
6. After forgetting the speech, “the judges all gave me/Liu Yang understanding smiles.”
7. When presenting the research topic, “I/Liu Yang was listened to and understood attentively.”
8. On the bus, I/Liu Yang forgot to bring money. “Many people helped me/him out.”
9. When I/Liu Yang encountered difficulties in my studies, “many of my/his/her classmates were willing to help me/him/her.”
10. When inviting friends to go out for fun, “I/Liu Yang always get an affirmative reply.”
11. When meeting a stranger, “I/Liu Yang received a polite and warm smile.”
12. I/Liu Yang handed out the leaflets to the passers-by. “The passers-by kindly accepted the leaflets.”
13. When I/Liu Yang express(es) negative emotions, “my/his/her friends offer me/him/her comfort and guidance.”
14. When I/Liu Yang accidentally fell down, “someone kindly helped me/him/her up and asked how was it going.”
15. When I/Liu Yang was paying at the supermarket but couldn’t find my/his/her wallet, “the cashier smiled at me/him/her and patiently waited.”
16. When I/Liu Yang showed confusion about something unfamiliar, “the other person responded calmly and provided an explanation”.
17. “The students think that I/Liu Yang am/is an easy-going person and they like me/him/her.”
18. When I/Liu Yang gave a gift to my/his/her friends, “they showed great joy.”
19. After presenting my/his/her ideas at the meeting, “everyone nodded in approval and encouraged me//Liu Yang to elaborate further.”
20. I/Liu Yang excitedly recounted the rare experiences had encountered, “friends listened attentively and all expressed their feelings together.”
21. I/Liu Yang didn’t perform well in this exam. “The teacher not only didn’t criticize me/Liu Yang, but also encouraged me/him/her not to give up.”
22. When I/Liu Yang was chatting with strangers, “I/Liu Yang received a friendly smile.”
23. When facing a difficult problem, I/Liu Yang proposed a solution approach, which “was appreciated and adopted by the partners”.
24. I/Liu Yang failed to complete the task. “My/His/Her teacher not only didn’t scold me/Liu Yang, but also asked me/him/her to tell her if I/Liu Yang had any problems.”
25. I/Liu Yang was not feeling well and was unable to perform my/his/her duties. “My/His/Her colleagues kindly came forward to inquire about my/his/her condition and assured me/him/her to rest.”
26. I/Liu Yang got a new hairstyle that was quite different from the previous one. “My/His/Her friends said it suits me/him/her well and even asked me/him/her to recommend a hairdresser.”
27. Following the map’s directions but still failing to find the place, “a stranger accompanied me/Liu Yang and we/they finally reached the destination together.”

**Negative social scenarios**

1. I/Liu Yang suddenly went out of tune during the singing competition, “causing the whole audience to burst into laughter.”
2. During my/his/her public speech, I/Liu Yang suddenly forgot the words. “The judges all looked sternly at me/him/her.”
3. After I/Liu Yang finished the speech contest, “not many of the audience liked my/his/her performance.”
4. During the class group discussion, “no one was willing to discuss with me/Liu Yang”.
5. “Not many people like me/Liu Yang.”
6. I/Liu Yang summoned up my courage and invited the person I/he/she liked to go out for fun, “but was rejected.”
7. When presenting the research topic, “no one listened to what I/Liu Yang was saying.”
8. When I/Liu Yang talked to strangers, “the other person had a stern expression on their face.”
9. I/Liu Yang actively proposed solutions to the problem, saying, but “the partner did not provide any feedback.”
10. I/Liu Yang politely handed out the leaflets to the passers-by, “but the passers-by ignored me/him/her and didn’t take the leaflets.”
11. When I/Liu Yang accidentally fell down, “the onlookers were pointing at me/him/her and laughing.”
12. When I/Liu Yang was paying at the supermarket but couldn’t find my/his/her wallet, “the cashier had an impatient expression and spoke in an unpleasant tone.”
13. When I/Liu Yang showed confusion about something unfamiliar, “the others wore mocking expressions and took pleasure in it.”
14. When I/Liu Yang passed by my classmates, “they would suddenly stop chatting and deliberately avoid me/him/her.”
15. When I/Liu Yang sought help from my/his/her friends, “they ignored me/him/her completely.”
16. After presenting my/his/her ideas at the meeting, “the meeting room was silent and no one gave me/Liu Yang any feedback.”
17. I/Liu Yang excitedly recounted the unusual things I/he/she had encountered on the way to work, “my/his/her friends just nodded and then changed the topic.”
18. I/Liu Yang couldn’t find the place and asked a passer-by for help. “Before I/Liu Yang could finish my/his/her question, the passer-by just walked away directly.”
19. I/Liu Yang happily invited friends to my/his/her birthday party, “but none of them were willing to come.”
20. The classroom for the specialized course was changed temporarily. The teacher informed everyone to pass on the message, “but no one told me about this.”
21. When I/Liu Yang forgot to bring money on the bus, “Everyone looked at me/him/her in a reproachful manner. Nobody helped me/him/her. I/Liu Yang had no choice but to get off the bus.”
22. When facing difficulties in study, “no one is willing to help me/Liu Yang.”
23. I/Liu Yang didn’t do well in the final exam for some reasons, “I/Liu Yang was severely criticized by the teacher.”
24. I/Liu Yang wanted to offer help to a stranger, “but the stranger showed no reaction.”
25. I/Liu Yang failed to complete task, “the teacher severely criticized me/him/her.”
26. When I/Liu Yang expressed negative emotions, my/his/her friends ignored it, didn’t care about it and take any action.
27. I/Liu Yang was not in good health and unable to perform job duties. “However, my/his/her colleagues all claimed they were busy and unwilling to hand over tasks.”

*MRI Data Acquisition*

Neuroimaging data were acquired using a 3T Siemens Prisma scanner (Siemens Healthineers, Erlangen, Germany) equipped with a 32-channel head coil at the University of Macau. Whole-brain functional images were obtained using a multiband-accelerated EPI sequence (MB factor = 5) with the following parameters: TR = 1000 ms, TE = 30 ms, flip angle = 90°, voxel size = 2×2×2 mm³, matrix size = 96×96, 84 axial slices with interleaved acquisition, phase encoding direction = posterior-anterior (j-), bandwidth = 2365 Hz/pixel, and echo spacing = 0.55 ms. The partial Fourier factor was set to 7/8 with in-plane parallel imaging (GRAPPA factor = 2). High-resolution T1-weighted anatomical images were acquired using an MPRAGE sequence: TR = 2300 ms, TE = 2.26 ms, TI = 900 ms, flip angle = 8°, isotropic voxel size = 1×1×1 mm³, matrix size = 256×256, and sagittal acquisition orientation. Parallel imaging was employed with an acceleration factor of 3 in the phase-encoding direction.

*MRI Data Preprocessing*

Functional MRI data preprocessing was conducted using SPM12 (Wellcome Department of Imaging Neuroscience, London, UK) within the MATLAB R2021b environment. The initial pipeline involved the realignment of functional images to the mean volume via a rigid-body transformation to mitigate head motion effects. Subsequently, the mean functional image was coregistered to each participant’s T1-weighted anatomical scan, followed by the tissue segmentation of the structural images. These images were then spatially normalized to the Montreal Neurological Institute (MNI) standard space using a nonlinear registration algorithm and resampled into isotropic voxels. Notably, spatial smoothing was applied selectively based on the analysis type. For the univariate parametric GLM, images were smoothed with a 6-mm FWHM Gaussian kernel to optimize the signal-to-noise ratio. In contrast, the MVPA utilized unsmoothed data to preserve fine-grained spatial patterns.

*MVPA Decoding*

To control for potential confounding factors such as scanner signal drift and within-session fluctuations in participant states (e.g., arousal or fatigue), we implemented a temporally-matched cross-run validation scheme. Specifically, for participants in the 4-run condition, a 2-fold split-half procedure was employed: the first pair of “Self” and “Other” runs was used to train the classifier, which was then tested on the second pair, and vice versa. For participants in the 6-run condition, we utilized a 3-fold validation scheme. In each fold, one temporally-aligned pair of “Self” and “Other” runs (e.g., Self Run 1 and Other Run 1) was reserved for testing, while the remaining two pairs were used for training. This approach ensured that the training and testing sets always maintained a strictly balanced 1:1 category ratio and that each fold compared neural representations captured at similar temporal stages of the experimental session.

Decoding was performed using the CoSMoMVPA toolbox (Oosterhof et al., 2016) implemented in MATLAB. To strictly control for potential confounding factors such as scanner signal drift and within-session fluctuations in participant states (e.g., arousal or fatigue), we implemented a temporally-matched cross-run validation scheme, ensuring that the training and testing sets always maintained a strictly balanced 1:1 category ratio (see Supplemental Methods).

Within this framework, a Linear Discriminant Analysis (LDA) classifier was trained to distinguish between the neural patterns of the target pairs, such as Self-Negative versus Other-Negative trials. To ensure the results were not biased by potential imbalances in trial counts, we balanced the partitions by matching the minimum number of trials across classes within each cross-validation fold. Group-level decoding accuracy was tested against the theoretical chance level (0.5) using a one-sample *t*-test (one-tailed). The one-tailed test was chosen because decoding accuracy is theoretically meaningful only above the chance level. P-values were then corrected for multiple comparisons across all ROIs using the False Discovery Rate (FDR) method.

*Parametric Modulation Analysis*

Individual subject analyses were conducted on spatially smoothed volumes (6-mm FWHM) using a GLM framework in SPM12. For each participant, the design matrix incorporated four primary experimental conditions: self-positive, self-negative, other- positive, and other-negative. Each condition was modeled as a boxcar function with a 6 s duration, convolved with the canonical HRF. To identify brain regions tracking subjective evaluations, subjective ratings on perceived social acceptance, anxiety, and future likelihood prediction were included as linear parametric modulators for each of the four conditions. Crucially, all parametric modulators were mean-centered within each run to ensure that the resulting beta weights represented the neural sensitivity to trial by trial fluctuations rather than mean activation levels. We utilized the default serial orthogonalization setting in SPM12, entering the parametric modulators in the following order: (1) social acceptance, (2) perceived anxiety, and (3) future likelihood prediction. This hierarchical approach allowed us to identify neural activity specifically associated with likelihood prediction that could not be accounted for by affective or evaluative factors (i.e., social acceptance and perceived anxiety). Six motion parameters derived from realignment were included as nuisance covariates. High-pass filtering (cutoff = 128 s) was applied to remove low-frequency drifts. At the group level, significant brain regions were localized using MNI coordinates (mm) and the AAL3 atlas.

*Mediation Analysis*

The mediation analysis was implemented via the lavaan package in R (Rosseel, 2012). We constructed a path model where SA (LSAS total scores) served as the independent variable, the representational distinction in the left pgACC (decoding accuracy) was the mediator, and the neural sensitivity to social prediction in the precuneus (parametric modulation effect) was the outcome variable.

Specifically, the model estimated the effect of SA on the left pgACC decoding accuracy (Path *a*), and the subsequent effect of the left pgACC on the precuneus response while controlling for SA (Path *b*). The indirect effect was calculated as the product of these coefficients (*a* × *b*), while the direct effect (*c’*) represented the relationship between SA and the precuneus response. To ensure the robustness of the results, we employed non-parametric bootstrapping with 5,000 iterations to calculate 95% bias-corrected confidence intervals (CIs). An indirect effect was considered statistically significant if the 95% CI did not encompass zero. All variables were standardized prior to model fitting to provide standardized path coefficients.

**Supplementary fMRI Results**

*Self versus Other Referential Processing*

To investigate the neural substrates of self-other differentiation during social processing, we conducted pairwise *t*-tests on the parameter estimates derived from the univariate GLM. During the processing of positive social outcomes, significantly stronger activations were observed for Self > Other in the medial prefrontal cortex (mPFC), anterior cingulate cortex (ACC), and caudate (Table S7; Cluster-level *p*_FWE_ < .05, Peak-level *p* < .001). In contrast, no regions showed significantly greater activation for Other > Self in the positive condition. Furthermore, no significant clusters were identified for the self-other contrast during the processing of negative social outcomes at the specified threshold. These findings suggest that the neural differentiation between self and other is specifically pronounced during the processing of rewarding social information, aligning well with the established “self-positivity bias” represented in the mPFC (Fields et al., 2019; Korn et al., 2012).

*Modulation Effect of Social Anxiety on Self versus Other Referential Processing*

We further conducted an exploratory analysis to investigate the modulation effect of social anxiety on self-/ other- referential processing. To achieve this, we conducted two whole-brain univariate second-level regression analyses, separately using the contrast maps of “Self versus Other” Positive and Negative conditions as the dependent variable, and the Liebowitz Social Anxiety Scale (LSAS) total score as a subject-level covariate. Analyses were implemented in SPM12. Due to the exploratory nature of this whole-brain search, we applied a lenient threshold of *p*_uncorrected_ < 0.005 at the peak-level, with a cluster size *k*_E_ > 8 voxels.

The results revealed that in the positive condition (Table S8), higher social anxiety was associated with increased neural activation difference (self > other) in a social cognitive brain regions, including the right middle frontal gyrus and the left pregenual anterior cingulate cortex (pgACC). This finding provides converging evidence for our primary ROI-based decoding accuracy regression model, which identified the decoding accuracy in the pgACC as a strongest predictor of social anxiety.

In the negative condition (Table S9), it revealed that social anxiety was associated with increased self versus other activation difference in social cognitive and affective brain regions, including the left middle frontal gyrus and the insula.
